# Mechanisms of resistance to VHL loss-induced genetic and pharmacological vulnerabilities

**DOI:** 10.1101/2025.06.14.659649

**Authors:** Jianfeng Ge, Shoko Hirosue, Saroor A. Patel, Ludovic Wesolowski, Anna Dyas, Cissy Yong, Leticia Castillon, Sanne de Haan, Jarno Drost, Grant D. Stewart, Anna C. Obenauf, Daniel Muñoz-Espín, Sakari Vanharanta

## Abstract

The von Hippel-Lindau tumor suppressor (VHL) is a component of a ubiquitin ligase complex that normally controls cellular responses to hypoxia. Endogenous VHL is also utilized by proteolysis-targeting chimera (PROTAC) protein degraders, a promising class of anti-cancer agents. VHL is broadly essential for cell proliferation, yet it is a key tumor suppressor in renal cell carcinoma. To understand the functional consequences of *VHL* loss, and to identify targeted approaches for the elimination of *VHL* null cells, we have used genome-wide CRISPR-Cas9 screening in human renal epithelial cells. We find that, upon VHL loss, the HIF1A/ARNT complex is the central inhibitor of cellular fitness, suppressing mitochondrial respiration, and that VHL null cells show HIF1A-dependent molecular vulnerabilities that can be targeted pharmacologically. Combined VHL/HIF1A inactivation in breast and esophageal cancer cells can also provide resistance to ARV-771, a VHL-based bromodomain degrader that has anti-cancer activity. HIF1A stabilization can thus provide opportunities for early intervention in neoplastic *VHL* clones, and the VHL-HIF1A axis may be relevant for the development of resistance to the emerging class of PROTAC-based cancer therapies.

## INTRODUCTION

Heterozygous germline mutations in the von Hippel-Lindau tumor suppressor (*VHL*) lead to a tumor predisposition syndrome that is characterized by the development of phaeochromocytomas, renal cell carcinomas, hemangioblastomas and pancreatic neuroendocrine tumors ^1^. Biallelic inactivation of *VHL* is also the tumor-initiating genetic event in ∼90% of sporadic clear cell renal cell carcinomas (ccRCCs) ^2^. The VHL protein functions as a substrate recognition subunit of an E3 ubiquitin ligase complex, the best-characterized substrates of which are the hypoxia-inducible factors HIF1A and HIF2A (HIFA). HIFA bind VHL when two of their conserved proline residues are hydroxylated ^1^. The HIFA prolyl hydroxylases EGLN1-3 require oxygen as a co-substrate, making them active only in normoxic conditions. Under hypoxia or in the absence of functional VHL, HIFA accumulates and dimerizes with HIF1B/ARNT, forming a helix-loop-helix transcription factor ^1^.

The endogenous ubiquitin ligase target recognition function of VHL is exploited by the VHL-dependent PROTACs, small molecules that can be engineered to degrade specific endogenous proteins through the E3 ubiquitin ligase pathway ^3^. These compounds serve as molecular bridges between a target protein of interest and one of a few E3 ligases, such as VHL, which directs them for proteasomal degradation. For example, ARV-771, a bromodomain degrader, and AU-15330, a SMARCA2/4 degrader, are dependent on endogenous VHL and they have shown anti-tumor activity in experimental cancer models ^4,5^.

Even though the cellular consequences of VHL inactivation are clinically relevant in several contexts, they remain incompletely understood. For example, acute VHL loss is detrimental to cell proliferation ^6^, yet *VHL* is a tumor suppressor in renal cancer ^2,6,7^. As the development of renal cancer from *VHL* mutant renal epithelial cells can take decades ^8^, VHL loss-induced vulnerabilities could be exploited for early cancer intervention strategies, especially in high-risk individuals, such as *VHL* mutation carriers. Bi-allelic *VHL* inactivation in the germline can also lead to a systemic metabolic disorder ^9^. Such patients could benefit from therapies that limit the negative fitness effects caused by *VHL* inactivation. Finally, the role of endogenous VHL as a key player in PROTAC-induced anti-cancer effects ^3^ suggests that understanding the consequences of VHL loss could be important for understanding and reducing the likelihood of PROTAC-resistance in cancer.

Using experimental cell line systems and large-scale CRISPR-Cas9 functional screening we set out to investigate the mechanisms of VHL loss-induced proliferative suppression. We find that the HIF1A/ARNT complex is the central mediator of reduced fitness following VHL loss, and that VHL null cells display HIF1A-dependent genetic vulnerabilities that can be pharmacologically targeted. Moreover, we demonstrate that combined loss of VHL and HIF1A can provide a fitness advantage to human cancer cells under treatment with a VHL-dependent PROTAC. These results shed light on the functional consequences of VHL loss at the genome-wide scale and provide a proof of principle that VHL loss-dependent phenotypes can be pharmacologically targeted.

## RESULTS

### HIF1A/ARNT-mediated proliferative suppression upon *VHL* inactivation

To investigate the effects of VHL loss in *VHL* wildtype human cells we used CRISPR-Cas9 to mutate *VHL* in the HK2 human renal epithelial cells, an immortalized cell line derived from human renal epithelial cells. This led to strong inhibition of proliferation and an altered cellular morphology (**Fig. S1A-C**). Reintroduction of sgRNA-insensitive *VHL* cDNA completely rescued the proliferation defect caused by CRISPR-Cas9-mediated *VHL* inactivation, confirming the specificity of the approach (**Fig. S1D-E**). Unlike previously reported in mouse fibroblasts ^7^, reduced oxygen level did not rescue the effects of *VHL* loss (**Fig. S1F**), even though it reduced the proliferative capacity of *VHL* wildtype cells in longer-term assays (**Fig. S1G)**. Analysis of the large-scale Cancer Dependency Map data set revealed that apart from ccRCC cells, most of which already carry biallelic mutations in the *VHL* gene, cancer cell lines across different non-renal lineages were universally sensitive to VHL inactivation (**Fig. 1A**). Inactivation of *VHL* also reduced the size of human renal epithelial organoids, but the cells still formed similar structures as wildtype cells (**Fig. 1B** and **Fig. S1H**). We generated a genetic GFP-dependent HIFA reporter to confirm HIF activity, a predicted downstream effect of VHL loss, in the VHL-engineered organoids, while wildtype organoids remained GFP negative (**Fig. 1C**).

**Fig. 1.**
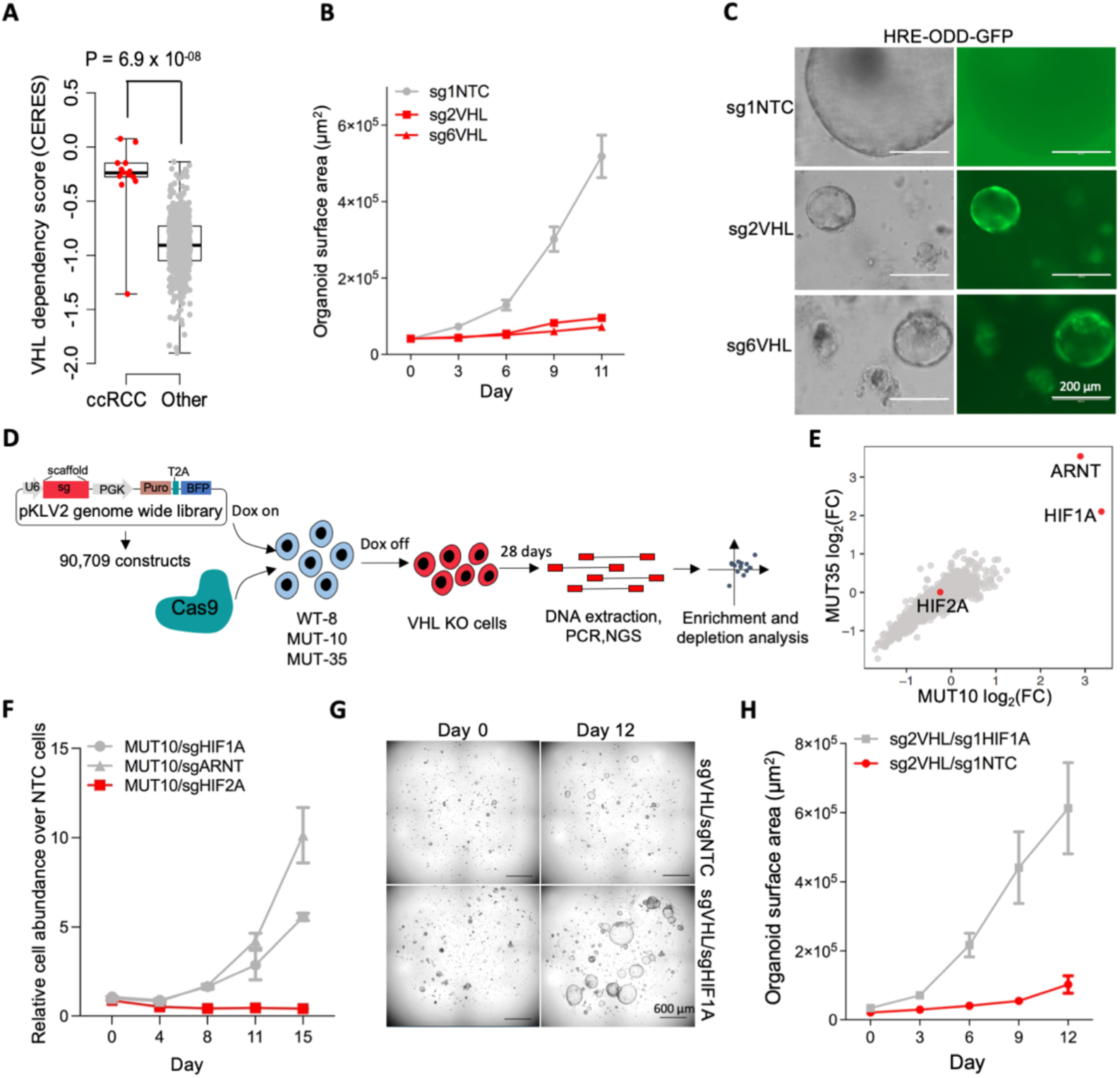
Universal inhibition of cell proliferation upon *VHL* inactivation. **A.** VHL dependency score in ccRCC cell lines and other pan cancer lines (Mean and S.E.M). **B.** Quantification of normal renal organoid growth (EMKC016) with and without VHL inactivation, N=32 random growing organoids per condition and time point (Mean and S.E.M). **C.** Activity of a hypoxia reporter (HRE-ODD-GFP) in normal renal organoids (EMKC016) with and without VHL deletion as determined by fluorescence microscopy. **D.** Schematic of the pooled CRISPR-Cas9 screening strategy. **E.** Gene level CRISPR-Cas9-based loss of function screening data. Beta scores showing change in sgRNA construct abundance in VHL mutant cells. N=2 replicates per condition. **F.** CRISPR-Cas9 based competition assay in VHL mutant MUT10 cells. HIF1A, HIF2A and ARNT mutant cells competed against cells transduced with non-targeting control constructs (NTC). Two sgRNAs per gene combined, N=3 technical replicates per condition (Mean and S.E.M). **G.** VHL mutant human renal epithelial organoids with or without HIF1A inactivation at different time points. **H.** Quantification of organoid growth from panel G over time, N=13 random growing organoids per condition and time point (Mean and S.E.M).

To understand the consequences of VHL loss, we performed a genome-wide CRISPR-Cas9 loss-of-function screen in HK2 cells. To avoid the possibility of cells adapting to VHL inactivation during propagation before the screen started and to obtain enough VHL null cells despite their reduced proliferative capacity, we engineered cell clones in which VHL expression could be regulated by doxycycline (dox), and endogenous *VHL* was either intact (WT8 control cells) or knocked out (MUT10 and MUT35) (**Fig. S2A**). WT8 cells proliferated well regardless of dox, but MUT10 and MUT35 cells proliferated significantly better when dox, and consequently VHL, was present (**Fig. S2B-D**). VHL expression was also associated with HIF1A and HIF2A degradation, as expected (**Fig. S2E**). Cas9 editing efficiency was confirmed in all three clones (**Fig. S2F**) before proceeding to the genome-wide screen (**Fig. 1D**). The sgRNA distribution of the two *VHL* mutant clones showed strong correlation at the endpoint (**Fig. S3A**). When comparing sgRNA representation in VHL null cells at the end of the screen and pre-dox withdrawal, constructs targeting HIF1A, a known target of VHL, and its dimerization partner ARNT were clearly the most significantly enriched, with little evidence of consistent enrichment of constructs targeting other genes (**Fig. 1E**). Consistenly, all five sgRNAs targeting HIF1A and ARNT showed enrichment (**Fig. S3B-C**). *VHL* null cells were sensitive to the inhibition of several pathways, such as glycolysis, oxidative phosphorylation, MTORC1 signaling, DNA repair, Myc and E2F targets and the G2M checkpoint (**Fig. S3D)**, pathways that are typically important for proliferating cells, indicating that these cells, despite their reduced proliferative capacity in comparison to *VHL* wildtype cells, performed as expected in the pooled screen. Fluorescence-assisted cell sorting-based competition assays confirmed that HIF1A and ARNT inactivation, but not HIF2A inactivation, rescued the proliferation defect caused by *VHL* loss (**Fig. 1F** and **Fig. S3E-H**). HIF1A and ARNT inactivation also restored the morphological appearance of *VHL* null cells (**Fig. S3I)**. Finally, HIF1A inactivation could rescue proliferation in *VHL* mutant human renal epithelial organoids (**Fig. 1G-H**), demonstrating the relevance of our observation in primary human cells. Overall, these results suggest that the HIF1A/ARNT complex is a central mediator of VHL loss-induced proliferative suppression.

### HIF1A-dependent mitochondrial inhibition in *VHL* null cells

To understand how gene expression was affected by *VHL* inactivation, we performed transcriptomic analysis by RNA-seq on MUT10, MUT35 and WT8 cells with and without dox withdrawal. As expected, WT8 cells showed little change in gene expression (**Fig. S4A** and **Supplementary Table 1**). However, combined analysis of MUT10 and MUT35 cells revealed 220 genes significantly upregulated and 63 genes significantly downregulated upon dox withdrawal and consequent VHL depletion (**Fig. 2A, Fig. S4B-C** and **Supplementary Table 2-3**). Gene set enrichment analysis of the transcriptomic data showed that genes in categories such as hypoxia, TNF-alpha signaling, glycolysis and epithelial-to-mesenchymal transition (EMT) were upregulated, and oxidative phosphorylation, MYC targets, G2M checkpoint and E2F targets were suppressed in VHL depleted cells (**Fig. 2B**). Although the expression of glycolysis genes is upregulated in *VHL* mutant cells, oxidative phosphorylation in mitochondria is the main source of ATP, suggesting that the proliferation defect of *VHL* mutant cells could be linked to suppressed oxidative phosphorylation (**Fig. 2C**). Oxygen consumption rate (OCR) in MUT10 and MUT35 cells in the absence of dox was reduced when compared to WT8 cells, with ATP production being ∼90% lower than in wildtype cells (**Fig. 2D-E**). In agreement with the cell proliferation data, HIF1A and ARNT inactivation rescued mitochondrial function and increased ATP levels in *VHL* null cells (**Fig. 2F-G**). As previously demonstrated ^10,11^, forced HIF1A expression also reduced proliferation and mitochondrial respiration in established ccRCC cell lines (**Fig. 2H-K, Fig. S4D-E)**. The HIF1A axis can thus reduce cell fitness in normal epithelial cells and advanced malignant cancer clones.

**Fig. 2.**
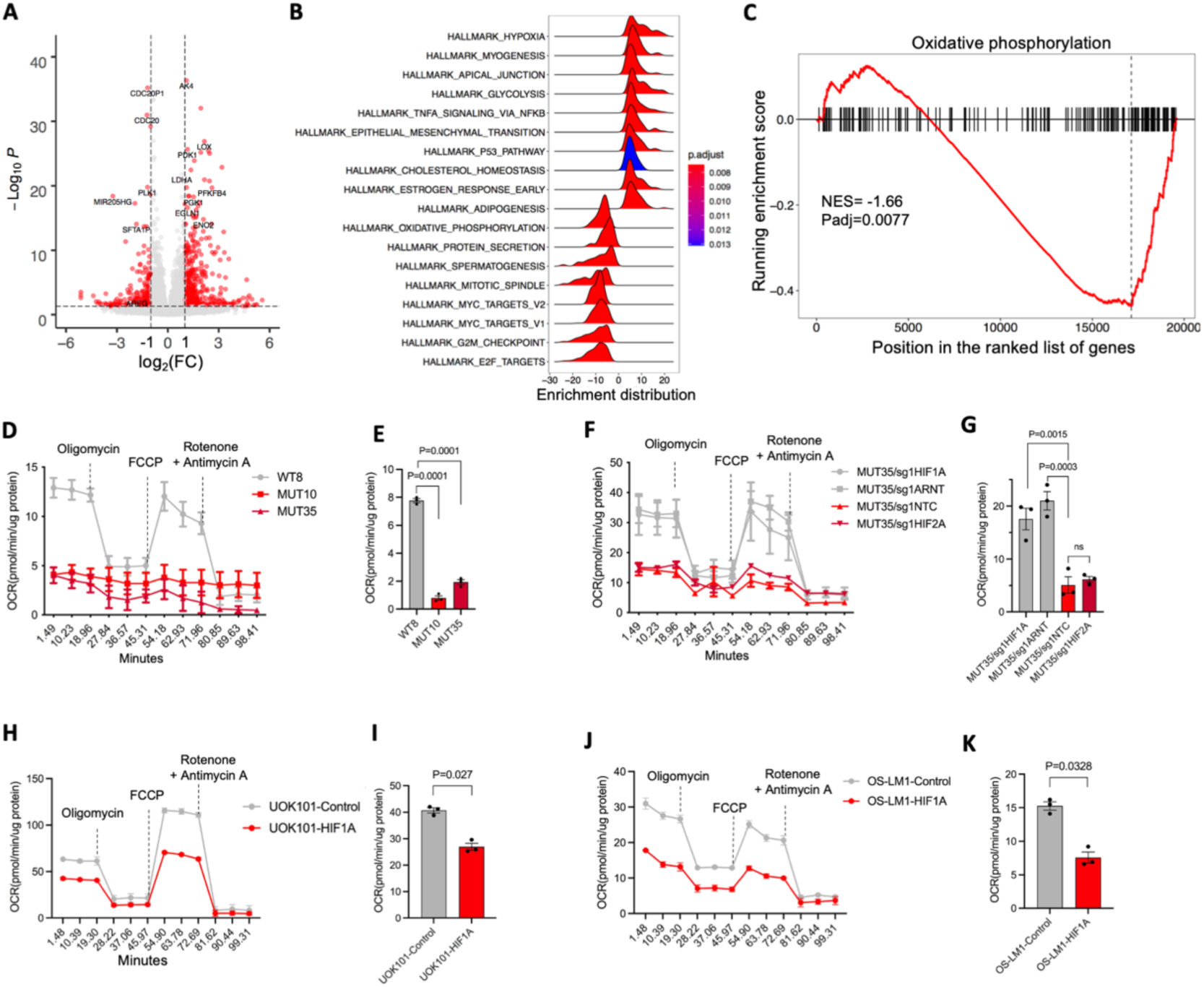
HIF1A stabilization induces mitochondrial inhibition and glycolysis dependency in VHL mutant cells. **A.** Differential gene expression analysis by RNA-seq. MUT10 and MUT35 cells compared to WT8 cells after dox withdrawal. P-value and fold-change determined by DESeq2. **B.** Gene set enrichment analysis (GSEA) comparing MUT10 and MUT35 cells to WT8 cells using the Cancer Hallmark gene set. **C.** Running GSEA enrichment score for the oxidative phosphorylation gene set. **D.** Oxygen consumption rate (OCR) in VHL WT and VHL MUT cells as measured by Seahorse analysis. N=3 per condition (Mean and S.E.M). **E.** Quantification of ATP production from mitochondria in VHL WT and VHL MUT cells. N=3 per condition. Dunnett’s multiple comparisons test (Mean and S.E.M). **F.** Seahorse assay to measure Oxygen consumption rate (OCR) in MUT35-NTC, MUT35-sgHIF1A, MUT35-sgHIF2A and MUT35-sgHIF1B cells. N=3 per condition (Mean and S.E.M). **G.** Quantification of ATP production from mitochondria in MUT35-NTC, MUT35-sgHIF1A, MUT35-sgHIF2A and MUT35-sgHIF1B cells. N=3 per condition. Sidak’s multiple comparisons test (Mean and S.E.M). **H.** Oxygen consumption rate (OCR) in UOK101 cells with and without HIF1A cDNA expression. N=3 replicates per condition (Mean and S.E.M) as determined by Seahorse analysis. **I.** ATP production in mitochondria in UOK101 cells with and without HIF1A cDNA expression. N=3 replicates per condition (Mean and S.E.M). Student’s t test. **J**. Oxygen consumption rate (OCR) in OS-LM1 cells with and without HIF1A cDNA expression. N=3 replicates per condition (Mean and S.E.M) as determined by Seahorse analysis. **K**. ATP production in mitochondria in OS-LM1 with and without HIF1A cDNA expression. N=3 replicates per condition (Mean and S.E.M). Student’s t test.

### HIF1A-dependent pharmacological vulnerabilities in *VHL* mutant cells

As HIF1A suppresses mitochondria, the main source of ATP, and as *VHL* null cells showed evidence of sensitivity to genes involved in glycolysis (**Fig. S3D**), we tested the possibility of using glycolysis inhibitors to specifically target VHL null cells. However, 2-deoxyglucose (2-DG) and AZD3965, inhibitors of hexokinase and monocarboxylate transporter 1/2, respectively, showed only a mild effect on either *VHL* mutant cells (**Fig. S5A-B)**. To explore the possibility that VHL loss resulted in other pharmacological vulnerabilities, we looked for potential druggable VHL loss-induced genetic dependencies in our CRISPR-Cas9 screen data. This identified FGFR1 and RAD51 as potential targets for which small molecule inhibitors were available (**Fig. S5C**). Pemigatinib and AZD4547 are kinase inhibitors with efficacy against FGFR1 ^12,13^. Both inhibited the proliferation of MUT10 and MUT35 cells when compared to WT8 cells (**Fig. 3A-B**). They also reduced ERK phosphorylation, a downstream effect of FGFR1 activation, more efficiently in MUT10 and MUT35 cells when compared to WT8 cells (**Fig. 3C-F**). B02, a RAD51 inhibitor ^14^, also inhibited the proliferation of MUT10 and MUT35 cells more efficiently when compared to WT8 cells (**Fig. S5D**). Importantly, however, cisplatin (CDDP), doxorubicin, imatinib and navitoclax showed broadly similar effects on *VHL* mutant and wildtype cells, demonstrating specificity of our observations (**Fig. S5E-H**). These results demonstrate that *VHL* mutations can induce pharmacological vulnerabilities with associated biochemical phenotypes in renal epithelial cells.

**Fig. 3.**
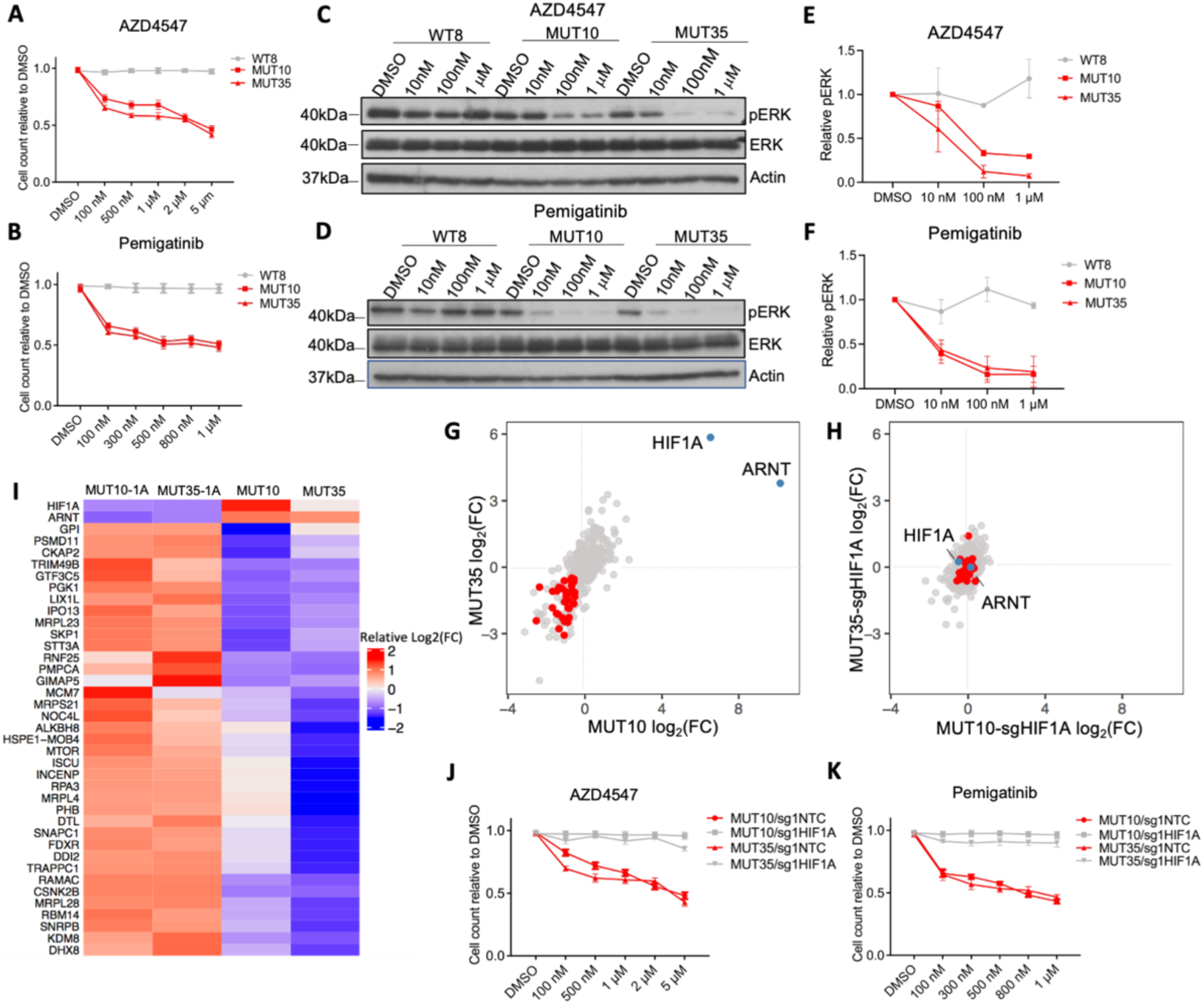
HIF1A dependent druggable genetic vulnerabilities in *VHL* mutant cells. **A,B.** Cell counts on day 5 relative to DMSO control group under AZD4547 (100 nM, 500 nM, 1 μM, 2 μM, 5 μM) and pemigatinib (100 nM, 300 nM, 500 nM, 800 nM,1 μM) treatments. N=4 per condition (Mean and SD). **C,D.** Western blot of pERK, ERK and beta-actin in WT8, MUT10 and MUT35 cells treated with AZD4547 (DMSO, 10 nM, 100 nM, 1 μM) and Pemigatinib (DMSO, 10 nM, 100 nM, 1 μM) **E,F.** Normalized pERK quantification in WT8, MUT10 and MUT35 cells treated with DMSO or AZD4547 (10 nM, 100 nM, 1 μM) or Pemigatinib (10 nM, 100 nM, 1 μM). N=2 per condition (Mean and S.E.M). **G-H** A targeted CRISPR-Cas9-based loss of function pooled screen in MUT10 and MUT35 cells (G), and double mutant MUT10-sgHIF1A and MUT35-sgHIF1A cells (H). HIF1A dependent gene dependencies are labelled in red. **I.** Heat map of 37 representative HIF1A dependent gene dependencies. Fold change in sgRNA abundance. **J-K.** Cell counts on day 5 relative to DMSO control group under AZD4547 (100 nM, 500 nM, 1 μM, 2 μM, 5 μM) and Pemigatinib (100 nM, 300 nM, 500 nM, 800 nM,1 μM) treatments. N=3-4 per condition (Mean and SD).

The central role of HIF1A in mediating VHL loss-induced proliferative suppression suggested that the VHL loss-induced genetic and pharmacological vulnerabilities could in general be HIF1A dependent. We tested this by a secondary pooled CRISPR-Cas9 screen that targeted genes the sgRNAs of which were depleted (94 genes) or enriched (250 genes) in *VHL* mutant cells in the original genome-wide screen (**Fig. S6A-B**). We also included constructs that targeted 30 genes frequently mutated in ccRCC and 22 non-essential control genes. The screen was performed using MUT10 and MUT35 cells together with the corresponding VHL-HIF1A double mutant cells MUT10-1A and MUT35-1A (**Fig. S6B**). Overall, the secondary screen validated the original results from the genome-wide screen in the *VHL* mutant cells (**Fig. 3G** and **S6C**). Interestingly, in the VHL-HIF1A double-mutant cells we observed little enrichment or depletion of sgRNAs, indicating that the genetic vulnerabilities identified in *VHL* mutant cells were HIF1A dependent in this context (**Fig. 3H-I**). Withdrawal of dox before lentiviral transduction of the sgRNA library also resulted in strong enrichment of HIF1A and ARNT constructs, indicating that the growth suppression caused by VHL loss and HIF1A stabilization was reversible (**Fig. S6D**).

To further test whether the pharmacological vulnerabilities were also HIF1A dependent we treated the *VHL* mutant and *VHL-HIF1A* double mutant cells with pemigatinib, AZD4547 and B02. In all cases *VHL-HIF1A* double mutant cells were significantly less sensitive when compared to *VHL* single mutant cells (**Fig. 3J-K** and **Fig. S6E**). The increased sensitivity of *VHL* null cells to ERK inhibition was also HIF1A dependent (**Fig. S6F-G)**. These data demonstrate that *VHL* mutant cells have pharmacological vulnerabilities that are largely HIF1A-dependent.

### Fitness advantage by VHL-HIF1A axis inactivation under PROTAC therapy

The E3 ubiquitin ligase activity of the VHL complex can be harnessed by PROTACs for targeted protein degradation in cells (**Fig. 4A**). While recent evidence from functional screening suggests that several E3 proteins can be utilized for proximity-induced protein degradation ^15^, the broadly essential role of VHL for cancer cell proliferation (**Fig. 1A**) makes it attractive for PROTAC applications in cancer. However, while VHL loss is expected to lead to PROTAC resistance, our results on renal epithelial cells suggested that a proliferation proficient PROTAC resistant phenotype would only emerge through combined loss of VHL and HIF1A/ARNT. We first tested the effect of ARV-771, a VHL-dependent bromodomain degrader that has previously shown anti-cancer activity ^5^, on a panel of cancer cell lines and the renal epithelial cell line HK2. All but HK2 cells were sensitive to ARV-771 (**Fig. 4B**). As predicted, OE33, MCF7, 1833-BoM, LNCaP and KBM7 cells became resistant to ARV-771 when VHL was inactivated (**Fig. 4C**). However, as HIF1A stabilization upon VHL suppression leads to reduced proliferative fitness, *VHL* single mutant cells were predicted not to be competitive even if they were drug resistant, whereas *VHL-HIF1A* double mutant cells were expected to be resistant and able to proliferate. To test this experimentally, we mixed wildtype (mCherry labelled, 40%), *VHL* mutant (GFP labelled, 55%) and *VHL-HIF1A* double mutant (BFP labelled, 5%) 1833-BoM, MCF7 and OE33 cells for triple competition assays under DMSO or ARV-711 treatment (**Fig. 4D** and **Fig. S7A-C**), and the proliferation of WT, *VHL* mutant and *VHL-HIF1A* double mutant OE33, 1833-BoM and MCF7 cells was also tested separately (**Fig. S7D-F**). In DMSO vehicle conditions for all cell lines, the wildtype cells had a proliferative advantage (**Fig. 4E-F** and **Fig. S7G-J**). However, when the cells were treated with ARV-771, the abundance of wildtype cells reduced quickly (**Fig. 4G-H** and **Fig. S7K-N**). The relative abundance of *VHL* mutant cells was briefly enriched, but eventually the *VHL-HIF1A* double mutant cells, even if present only as a small minority population at the start of the experiment, became the dominant population (**Fig. 4G-H** and **Fig. S7K-N**).

**Fig. 4.**
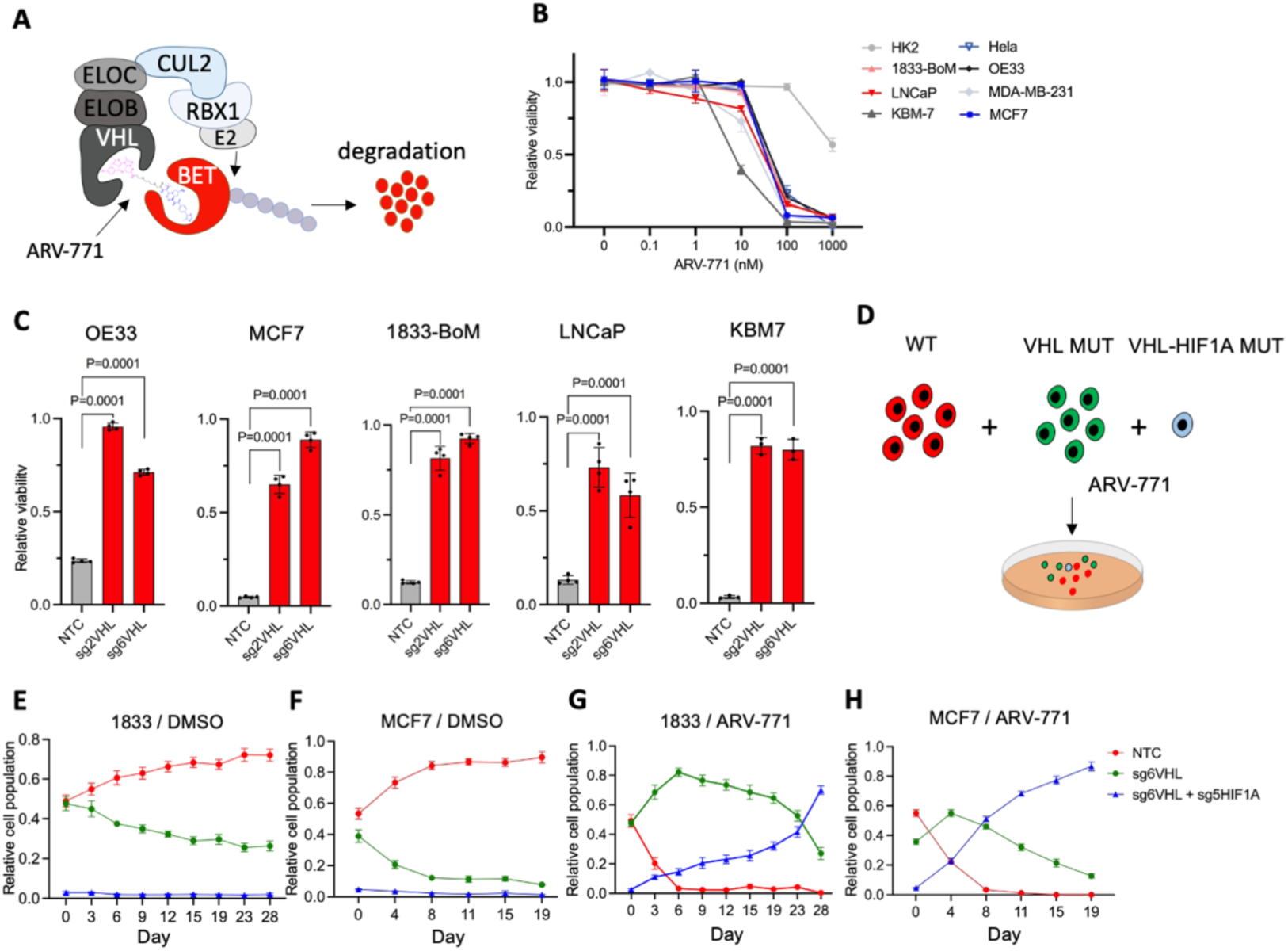
PROTAC resistance through HIF1A inactivation in cancer. **A.** Schematic of ARV-771 function as a PROTAC BET degrader. **B.** Relative viability of cells upon ARV-771 treatment (N=3 replicates per condition; error bar, SD; Dunnett’s multiple comparisons test). **C.** Relative cell viability upon ARV-771 treatment in wildtype and VHL mutant cells. ARV-771 concentrations as follows: OE33 (100 nM), MCF7 (1 μM), 1833-BoM (100 nM), LNCaP (100 nM), KBM7 (1 μM). N=3-4 replicates per condition (Mean and S.E.M). Dunnett’s multiple comparisons test. **D.** Schematic of long-term competition assay of WT (mCherry), VHL-MUT (GFP) and VHL-HIF1A double mutant (BFP) 1833-BoM, OE33 and MCF7 cells, treated with DMSO or ARV-771 (500nM for 1833-BoM, 400nM for OE33, 200 nM for MCF7). **E-H.** FACS-based quantification of the relative abundances of different cell populations in competition assays. WT (mCherry), VHL-MUT by VHLsg6 (GFP) and VHL-HIF1A double mutant by VHLsg6 and HIF1Asg5 (BFP). 1833-BoM cells: DMSO (E) and MCF7 cells: DMSO (F). 1833-BoM cells: 500nM ARV-771 (G) and MCF7 cells: 200 nM ARV-771 (H). N=3 for each condition and timepoint (Mean and SD).

To test whether the proliferative fitness advantage of *VHL-HIF1A* double mutant cells under ARV-771 treatment translateed into an advantage over *VHL* mutants cells also in vivo, we mixed wildtype (BFP), *VHL* mutant (GFP) and *VHL-HIF1A* double mutant (GFP+mCherry) MCF7 cells, and then performed an in vivo competition assay in a context of ARV-771 treatment for 21 days (**Fig. 5A**). The in vivo xenograft tumor assay confirmed that ARV-771 reduced tumor growth (**Fig. 5B**), and immunohistochemical staining showed less Ki67 and more cleaved caspase 3 in ARV-771 treated tumors (**Fig. 5C**). Importantly, based on flow cytometry analysis, the relative abundance of *VHL-HIF1A* double mutant cells was increased incomparison to *VHL* single mutant and wildtype cells in ARV-771 treated tumors but not in control tumors, indicating selective advantage of *VHL-HIF1A* double mutant cells under PROTAC therapy *in vivo* (**Fig. 5D**). These data demonstrate that inactivation of the VHL-HIF1A axis can cause functional resistance to VHL-dependent PROTACs in cancer cells.

**Fig. 5.**
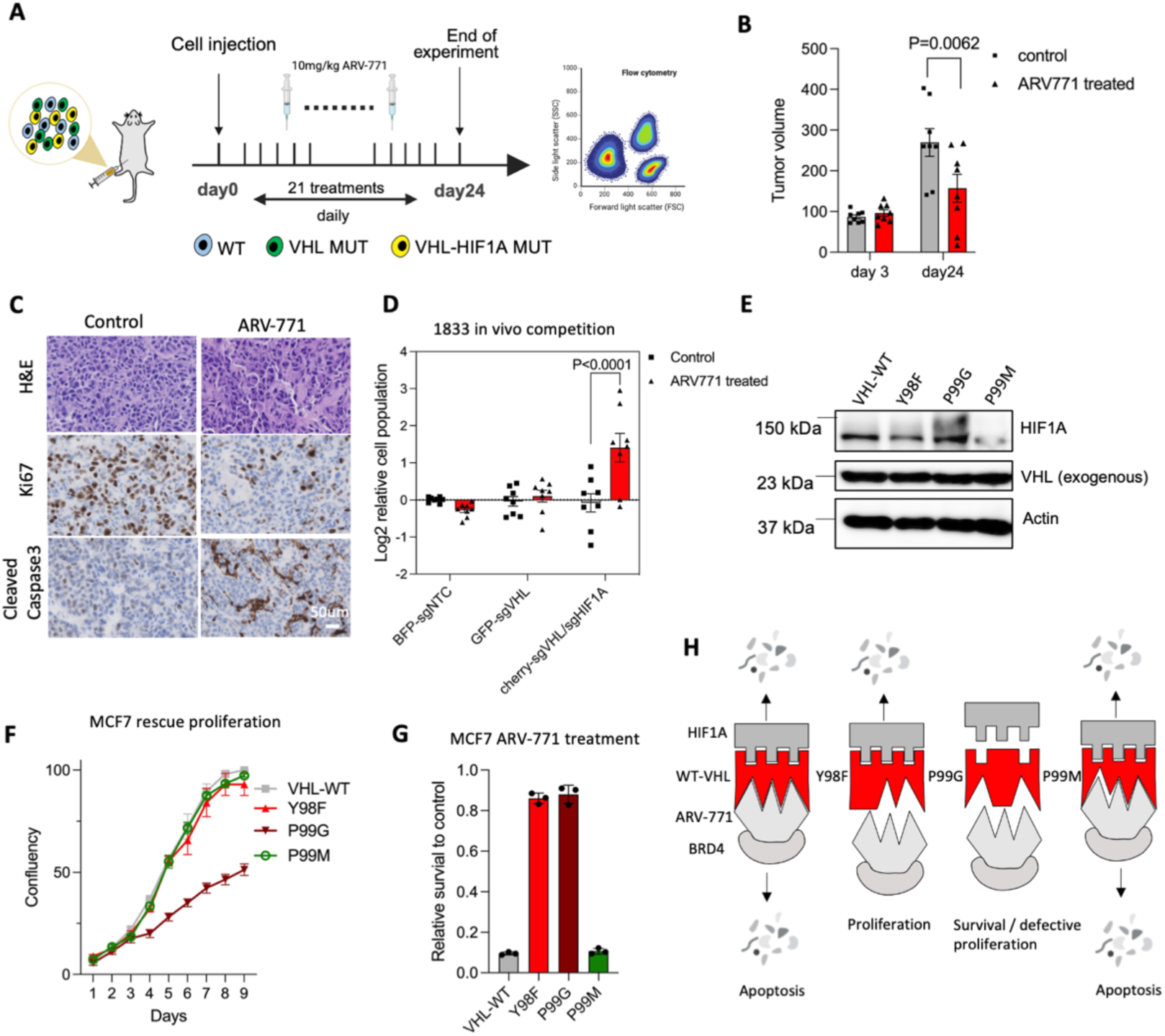
Uncoupling VHL functions in HIF1A regulation and PROTAC sensitivity. **A**. Schematic of in vivo competition assay of WT (BFP), VHL-MUT (GFP) and VHL-HIF1A double mutant (GFP+mCherry) MCF7 cells subcutaneously injected on nude mice, mice were daily treated with DMSO or 10mg/kg ARV-771 for 21 days. Tumors were dissociated for FACS analysis. **B**. Tumor size on day 3 and day 24 after inoculation of 2 million 1833-BoM cells into the flanks of athymic mice. N=8 tumors for each condition (Mean and S.E.M). Sidak’s multiple comparisons test. **C**. H&E, Ki67 and Cleaved Caspase 3 staining of tumors treated with DMSO and ARV-771. D. Log2 of relative cell population for each subpopulation of indicated treatment. N=8 tumors for each condition (Mean and S.E.M). Sidak’s multiple comparisons test. **E**. HIF1A and VHL western blot on VHL^WT^, VHL^Y98F^, VHL^P99G^ and VHL^P99M^ expressing VHL mutant MCF7. **F**. Incucytes proliferation of VHL-mutant MCF7 while expressing with VHL^WT^, VHL^Y98F^, VHL^P99G^ and VHL^P99M^. **G**. Relative cell viability upon 1 μM ARV-771 treatment in VHL^WT^, VHL^Y98F^, VHL^P99G^ and VHL^P99M^ expressing VHL-KO MCF7 cells. N=3. (Mean and S.E.M). **H**. Schematic of VHLY98F resistant to ARV-771 treatment and not activate HIF1A protein.

### Uncoupling VHL functions in HIF1A regulation and PROTAC sensitivity

Genotype-phenotype analyses on human *VHL* mutation carriers suggest that some *VHL* variants (type 2 mutations) predispose mainly to phaechromocytoma/paraganglioma while more severe alterations (type 1 mutations) are also associated with ccRCC ^16,17^. HIF stabilization is typically associated with type 1 mutations, while type 2 mutations may retain the capability to target HIFs for degradation ^17^. This raised the possibility that the ability of *VHL* to mediate PROTAC effects could also be uncoupled from its role in HIF1A regulation. Consequently, there could be *VHL* mutations that disrupt PROTAC activity but do not lead to HIF1A stabilization and reduced proliferative fitness. To test this, we examined data from recent *VHL* saturation mutagenesis screens ^22,24^ and identified point mutants of *VHL* residues Y98 and P99 as potential candidates. Using site-directed mutagenesis, we generated dox-inducible *VHL^Y98F^*, *VHL^P99G^*and *VHL^P99M^* expression constructs and transduced them into *VHL* knock-out MCF7 cells. Based on immunoblot analysis, *VHL^WT^*, *VHL^Y98F^* and *VHL^P99M^* were able to reduce HIF1A expression, whereas *VHL^P99G^* was not (**Fig. 5E**). In line with this, cells expressing *VHL^Y98F^* and *VHL^P99M^*were able to proliferate at the same rate as *VHL^WT^* cells, whereas *VHL^P99G^* cells proliferated significantly slower (**Fig. 5F**). However, while *VHL^WT^* and *VHL^P99M^* were sensitive to ARV-771, both *VHL^Y98F^* and *VHL^P99G^* showed only limited sensitivity (**Fig. 5G**). Thus, *VHL^Y98F^*was able to abrogate PROTAC sensitivity while maintaining proliferative capacity at the level of *VHL^WT^* cells (**Fig. 5H**). Altogether, our results show that HIF1A stabilization provides potential opportunities for early intervention in pre-neoplastic *VHL* clones. In addition, we propose that the VHL-HIF1A axis may be relevant for the development of proliferation-proficient therapy resistance to the emerging PROTAC-based anti-cancer strategies.

## DISCUSSION

The VHL-HIF1A pathway regulates metazoan responses to reduced oxygen availability, and the phenotypic sequelae of VHL inactivation are relevant in several clinical contexts ranging from carcinogenesis ^1^ and metabolic dysfunction ^9^ to novel cancer therapies ^3^. Using unbiased functional screening in experimental systems we have thus characterized the consequences of acute VHL loss, demonstrating that HIF1A activation in VHL null cells can lead to potentially actionable genetic vulnerabilities.

Our data suggest that HIF1A is the central mediator of VHL loss-induced negative fitness effects, with little involvement of HIF1A-independent pathways detected in our screen. This is in line with recent data on mouse embryonic fibroblasts ^18^, but earlier studies have also reported HIF1A-independent mechanisms through which VHL loss may inhibit cell proliferation ^6,7^. Interestingly, recent data suggest that in some cell types VHL loss-induced activation of HIF2A may be the critical effector of fitness loss, pointing at cell type-specific effects^19^. We show that VHL loss suppresses mitochondrial function, a result concordant with known effects of HIF1A on mitochondrial activity ^20,21^. Importantly, we demonstrate that HIF1A activation in VHL null cells can result in pharmacologically targetable genetic vulnerabilities, such as increased sensitivity to FGFR1 or RAD51 inhibition. These results provide a proof of principle that non-cancerous VHL-null cells can be susceptible to therapeutic intervention, complementing *VHL* synthetic lethal screens previously performed in already established *VHL* mutant renal cancer cell lines ^28,29^. For example, the cancer risk of individuals carrying heterozygous *VHL* mutations could be reduced even with incomplete early eradication of premalignant *VHL*-null clones. Expanding the concepts from our findings into in vivo models in which VHL-null cells could be analyzed in a non-proliferative state would be a critical next step. On the other hand, HIF1A-dependent suppression of mitochondrial activity in non-cancerous *VHL*-null cells could explain the systemic metabolic defects associated with biallelic germline mutations in *VHL* ^9^. Strategies to inhibit HIF1A activity could ameliorate the severe phenotype in such patients.

PROTACs are an emerging class of versatile molecules that can harness endogenous E3 ubiquitin ligase activity for specific degradation of target proteins of interest. Most PROTACs in clinical and pre-clinical development use CRBN or VHL as the endogenous E3 ubiquitin ligase proteins ^3^. Loss of the relevant E3 complex compromises PROTAC activity ^23^, but as VHL is broadly essential for cancer cell proliferation, VHL loss alone is unlikely to result in clinically relevant PROTAC resistance. Our data suggest that cell proliferation-proficient resistance to VHL-dependent PROTACs, intrinsically rare compared to resistance to CRBN-dependent PROTACS ^24^, can result from combined VHL and HIF1A inactivation, or through VHL mutations that retain the activity to target HIF1A for degradation. Reducing the probability of HIF1A inactivation in VHL deficient clones under therapy could improve long-term efficacy of PROTAC treatment. This could be achieved through the elimination of HIF1A-positive VHL-null cells before they lose HIF1A, possibly by targeting HIF1A-induced vulnerabilities as demonstrated by our data. Alternatively, the identification of synthetic lethal approaches for HIF1A null cells could be beneficial.

In conclusion, our data provide genome-wide functional insight into the molecular consequences of VHL loss in human cells. This revealed specific, mostly HIF1A-dependent genetic vulnerabilities in VHL null cells, suggesting the possibility that VHL loss-induced phenotypes could be exploited therapeutically. We also describe mechanisms that can provide cancer cells with proliferative fitness upon resistance to VHL-dependent PROTACs. This could aid the future development and clinical applicability of this promising novel class of anti-cancer agents.

## METHODS

### Cell lines

The HK2 cell line was obtained from C. Frezza (MRC Cancer Unit, Cambridge, UK). KBM7 cells were obtained from A. Obenauf (IMP, Vienna). OE33 cells were obtained from Rebecca Fitzgerald (MRC Cancer Unit, Cambridge, UK). LnCAP cells were obtained from Charlie Massie (Early Cancer Institute, Cambridge, UK). UOK101 cells were obtained from M. Linehan (NCI, Bethesda). OS-LM1, MDA-MB-231, 1833-BoM and 293T cells were obtained from J. Massagué (MSKCC, New York, USA). WT8, MUT10 and MUT35 are single cell clonal derivatives of HK2 cells with VHL mutation generated by CRISPR-Cas9 using VHLsg6, and they are available from the corresponding author upon reasonable request. VHL inactivation was validated by Sanger sequencing and Western blotting for VHL and HIF2A. The identity of the cell lines was confirmed by STR analysis. Cells were also confirmed to be mycoplasma free. HK2 and ccRCC cell lines were cultured in RPMI-1640 medium (Sigma) supplemented with 10% FBS, penicillin (100U/mL) and streptomycin (100μg/mL). All other cell lines were cultured in high glucose DMEM medium (Invitrogen) supplemented with 10% FBS, penicillin (100U/mL) and streptomycin (100μg/mL).

### Plasmids

psPAX2 and pMD2.G were gifts from Didier Trono (Addgene # 12260 and # 12259), lenti-cas9-blast was a gift from Feng Zhang (Addgene# 52962). pKLV2-U6gRNA5(BbsI)-PGKpuro2ABFP-W (Addgene#67974) was a gift from Kosuke Yusa. pKLV2-U6gRNA5(BbsI)-PGKhygro2ABFP, pKLV2-U6gRNA5(BbsI)-PGKpuro2AGFP and pKLV2-U6gRNA5(BbsI)-PGKpuro2AmCherry are derivatives of pKLV2-U6gRNA5(BbsI)-PGKpuro2ABFP-W. The dox-inducible shRNA expression plasmid LT3GEPIR was kindly gifted by J. Zuber (IMP, Vienna). sgRNA sequences are listed in **Supplementary Table 4**. HIF1A cDNA was amplified from HA-Clover-HIF-1alpha Wildtype (Addgene#163365). pLVX-puro (632164, Clonetech) was used to exogenously express the cDNA constructs. pLVX-HRE-ODD-GFP-hygro was generated by fusing GFP with HIF1A-ODD domain and cloned under a hypoxia response element modified minimal CMV promoter.

### Lentiviral production and transduction

HEK293T cells were transfected with a mixture of the lentiviral transfer plasmid containing genes of interest, psPAX2 and pMD2.G using PEI reagent (MW.25,000. Alfa Aesar). The supernatant containing the lentivirus was collected 72 hours post-transfection and filtered by a 0.45μM PVDF sterile filter (ELKAY). Cells were transduced with the lentiviral supernatant in the presence of 5 μg/mL polybrene (Millipore). Puromycin (4μg/mL), hygromycin (800μg/mL, InvivoGen) or blasticidin (10μg/mL, InvivoGen). Selection started 48 hours post-transduction.

### Pooled CRISPR-Cas9 screening

The lentiviral genome wide sgRNA library was produced using HEK293T cells as described above. A total of 450 million cells per condition were transduced with the lentiviral library at a low MOI (<0.3) to ensure that 95% of cells had a single sgRNA integration, resulting in 500 X sgRNA representation. Puromycin was added 48 hours post-infection. Dox was removed from culture media 7 days post-infection to stop transgene VHL expression. Pooled cells were allowed to proliferate for 4 weeks before harvested for DNA extraction. Genomic DNA was extracted using the QIAamp DNA Blood Maxi Kit (Qiagen Cat # 51192). All DNA was used for the amplification of the sgRNA cassette. Amplified product was purified by 1% agarose gel, quantified with the Qubit dsDNA HS assay kit (Thermo) and pooled in equimolar concentrations prior to Illumina sequencing on a HiSeq4000 instrument. Sequencing results were analyzed by the MAGeCK protocol ^25^.

### In vitro proliferation assays

For proliferation phenotype assays, all cells were plated on a 24-well plate in triplicates with a start confluency ranging from 10%-15%. Proliferation was measured by Incucyte 2020. For competition assays, control and target cells, which carried different fluorescent markers (BFP+/mCherry+/GFP+), were mixed and plated onto multi-well plates in triplicates. The percentage of each cell population was analyzed from day 0 and multiple time points throughout the assays by flow cytometry on LSR Fortessa (BD Biosciences). The following gating approach was used: FSC-A, FSC-W, SSC-A to select for live and single cells, and then BFP (383nm/445nm), mCherry (561nm/610nm), or GFP (488nm/510nm) channels for discriminating between the cell populations.

### Protein detection

Total protein was extracted from cell pellets using RIPA buffer (Sigma) containing protease K and phosphatase inhibitor cocktail (Sigma) according to the manufacturer’s protocol. Proteins were separated by SDS-PAGE, transferred onto PVDF membrane (Millipore) and blotted with VHL (BD Pharmingen, 565183, 1:500), HIF1A (Proteintech,20960-1-AP, 1:1000), HIF2A (Novus Biologicals, NB100-122, 1:1000), beta-actin (Sigma-Aldrich, A1978, 1:30000), p-ERK (Abcam, ab201015,1:1000), ERK (Abcam, ab184699, 1:2000) antibodies. Secondary antibodies were polyclonal goat anti-mouse IgG/HRP (Dako, P0447, 1:10000) and polyclonal goat anti-rabbit IgG/HRP conjugated (Dako, P0448,1:5000). Protein expression was quantified using Image J and normalized to beta-actin.

### RNA-seq

RNeasy Mini Kit (Qiagen 74104) was used for total RNA extraction on sub-confluent cells in four replicates according to the manufacturer’s protocols. The quality and concentration were assessed with the Agilent RNA Nano 6000 kit (Agilent 5067-1511) on Agilent Bioanalyzer 2100 instrument. RNA-seq libraries were prepared using the SENSE/CORALL mRNA-seq Library Prep Kit (Lexogen), 1μg of total RNA was used as the starting material following the manufacturer’s recommendations. The library size and quality of the final library products were assessed using Agilent High Sensitivity DNA Kit (Agilent 5067-4626). Library concentration was determined using the KAPA Library Quantification Kit (KR0405). Libraries were pooled in equimolar concentrations and subjected to Illumina sequencing on a HiSeq4000 instrument. Single clone RNA-seq sequencing reads were mapped to hg38 using RSEM and bowtie2. Differentially expressed genes were identified using DeSeq2 ^26^. Gene set enrichment analysis was performed using R packages ClusterProfiler ^27^ and Molecular Signature Database (MSigDB) Hallmarks gene set (Version 7.1.1).

### Seahorse experiments

To assess the oxygen consumption rate (OCR) 80,000 cells were seed in XF^e^ 24 well Cell Culture microplate in 100 μL normal RPMI night before experiment. The next day cells were washed once in PBS and the medium was replaced with 675 μL of XF RPMI Medium (Agilent Seahorse, 103576-100) supplemented with 25 mM glucose, 1mM pyruvate, 4mM glutamine. To eliminate residues of carbonic acid from medium, cells were incubated for at least 30 minutes at 37 °C with atmospheric CO2 incubator. OCR was assayed in a Seahorse XF-24 extracellular flux analyzer by the addition via ports A-C of 1μM oligomycin (port A), 1μM carbonyl cyanide-p-trifluoromethoxyphenylhydrazone (FCCP, port B), 1μM antimycin A (port C). Two or three measurement cycles of 2 min mix, 2 min wait, and 4 min measure were carried out at basal condition and after each injection. Each well was washed twice with 1 mL PBS and proteins were extracted with 50 μL of radioimmune precipitation assay (RIPA) lysis medium at room temperature end of the experiment. Protein concentration in each well was measured by a BCA assay according to the manufacturer’s instructions (Thermo). OCR values were normalized to total μg of proteins in each well.

### Human renal epithelial organoids

Human renal epithelial organoids were generated using a previously described method ^30^. Normal human kidney tissue was sampled with informed consent by a consultant uropathologist within 2 hours of nephrectomy under an ethical approval by the East of England - Cambridge Central Research Ethics Committee (19/EE/0161). The tissue was transported on ice in cold HBSS, dissected in PBS supplemented with penicillin (100U/mL)/streptomycin (100μg/mL) and further split into advanced DMEM/F12 (Gibco). Kidney tissue pieces were minced, washed in wash medium (advanced DMEM/F12 supplemented with 1X Glutamax, penicillin (100U/mL), streptomycin (100μg/mL) and 10mM Hepes) and resuspended in kidney organoid medium containing collagenase A (1mg/mL, Sigma) for 45 minutes at 37°C with shaking. The cells were washed, pelleted by centrifugation (5 minutes, 300 rcf, 4°C), resuspended in wash medium, passed through a 70-μm strainer, pelleted by centrifugation (5 minutes, 300 rcf, 4°C) and resuspended in 100μl wash medium. Single cells were seeded in 70% growth factor-reduced BME (R&D systems) and cast into 20ul droplets in a 12-well plate. After polymerization of the BME (30 minutes, 37°C), 1mL kidney organoid medium was added to each well. For passage, organoids were dissociated using 1mL TrypLE (Gibco) containing 10 μM Y-27632 per well (InvivoGen). TrypLE dissociation was stopped by adding 10 mL advanced DMEM/F12 and centrifuged at 300 rcf for 5 minutes, cells were reseeded in fresh 70% BME and topped with kidney organoid medium: ADMEM/F12 supplemented with 1.5% B27 supplement (Gibco,17504044), 40% Wnt3A conditioned medium, 10% RSPO-conditioned medium, EGF (50ng/mL, Proteintech), FGF-10 (100ng/mL, Proteintech), N-acetylcysteine (1.25mM/mL, Sigma), Y-27632 (10μM, Apebio) and primocin (100ng/mL, InvivoGen).

### Organoid lentiviral transduction

Lentiviral supernatant was collected and mixed with LentiX concentrator (TaKaRa) in a ratio of 3:1. The mixture was incubated for 30 minutes at 4°C, and centrifuged at 4°C (1500 rcf, 1 hour). Virus pellets were resuspended in renal organoid medium with polybrene (5 μg/ml). Organoids were dissociated into single cells and organoid pellets were resuspended in concentrated virus in 15 ml falcon tubes and incubated for 4 hours as previously described ^31^. The falcon tubes were then centrifuged at 600 rcf at 32°C for 1 hour before reseeding organoids in fresh 70% BME.

### Statistical analysis

Statistical analyses were performed either in R or GraphPad Prism. P-values lower than 0.05 were considered statistically significant. For correlation analyses, Person correlation coefficient was calculated. For drug treatment assays, two-way ANOVA with Sidak’s multiple comparisons test was used. For seahorse experiments, ATP quantification analysis, ordinary one-way ANOVA with Sidak’s multiple comparisons test was used.

### Animal studies

The animal experiment was performed in accordance with protocols approved by the Home Office (UK) and the University of Cambridge Animal Welfare and Ethical Review Body (P7EC604EE). For subcutaneous tumor growth assays, 3 x 10^6^ cells in 100µL of 1:1 PBS/Matrigel Matrix (BD) solution were injected into both flanks of 5–6-week-old athymic nude mice (Charles River Laboratories). Tumor growth was followed by caliper measurements. Tumor volume (V) was calculated using the equation V= (length x width^2^) x 0.5.

## Data availability

The RNA-seq data generated within this project have been uploaded into the Gene Expression Omnibus under the access code GSE213241. Other datasets generated during the current study are available from the corresponding author on reasonable request.

## ACKNOWLEDGEMENTS

We thank M. Linehan for the UOK101 cells. G.D.S. is supported by the Mark Foundation for Cancer Research and the Cancer Research UK Cambridge Centre (C9685/A25177). G.D.S. and the Human Research Tissue Bank were supported by the NIHR Cambridge Biomedical Research Centre (BRC-1215-20014). The views expressed are those of the author(s) and not necessarily those of the NIHR or the Department of Health and Social Care. S.H. received a PhD studentship from the Rosetrees Trust. This project has received funding from the European Union’s Horizon 2020 research and innovation program under the Marie Skłodowska–Curie grant agreement No 955951. This work was supported by the Medical Research Council (MC_UU_12022/7), Kidney Research UK (RP_033_20170303), the Sigrid Jusélius Foundation and the Academy of Finland (decision 338420). Munoz-Espin laboratory is supported by a CRUK Programme Foundation Award (C62187/A29760). JG is funded by a Darley/Sands Downing College Fellowship (G109261).

## AUTHOR CONTRIBUTIONS

J.G. designed and performed experiments, analyzed data and wrote the manuscript. S.H. performed computational analyses. S.A.P., L.W., A.D. and L.C. assisted with experiments and/or data analysis. C.Y. and G.D.S. provided human material. S.d.H. and J.D. assisted with the organoid experiments. A.C.O. designed experiments and analyzed results. D.M.E. supervised research. S.V. supervised the project, designed experiments, analyzed data and wrote the manuscript.

---

**Fig. S1.**
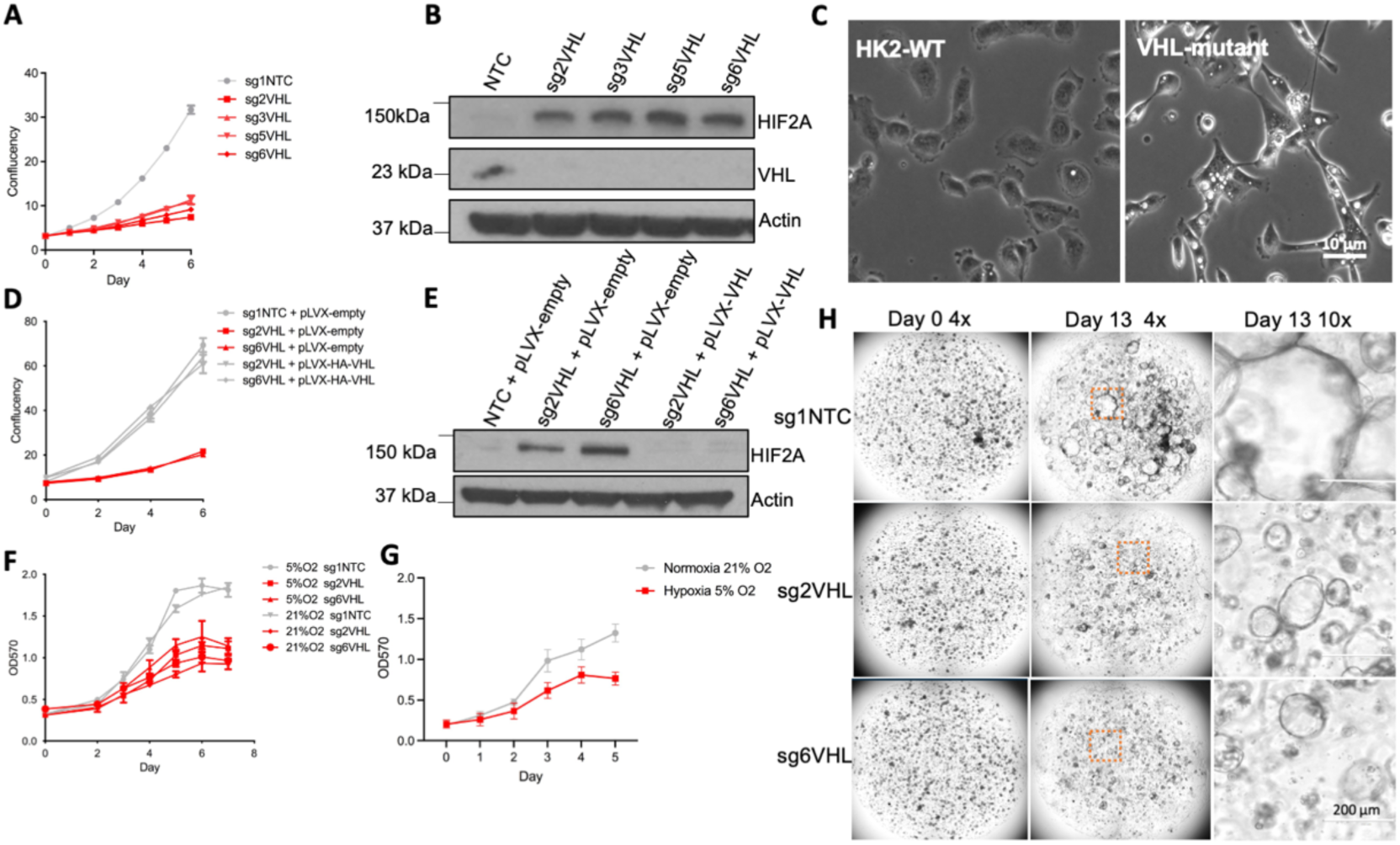
VHL inactivation inhibits cell proliferation. **A.** Proliferation of HK2 cells with or without VHL. N=2 per condition. (Mean and S.E.M). **B.** Western blot of HIF2A, VHL and Actin on HK2 cells with and without VHL. **C.** Morphology of HK2 cells with and without VHL. **D.** Proliferation of VHL mutant HK2 cells with and without VHL re-introduction. N=3 per condition. (Mean and S.E.M). **E.** Western blot of HIF2A on VHL mutant HK2 cells with and without VHL re-introduction. **F.** Proliferation of HK2 cells with and without VHL under 5% or 21% O_2_ culture conditions. N=2 per condition. (Mean and S.E.M). **G.** Representative images of human renal epithelial organoids with and without VHL. **H.** Human renal epithelial organoid proliferation under DMSO or DMOG (3mM) treatment.

**Fig. S2.**
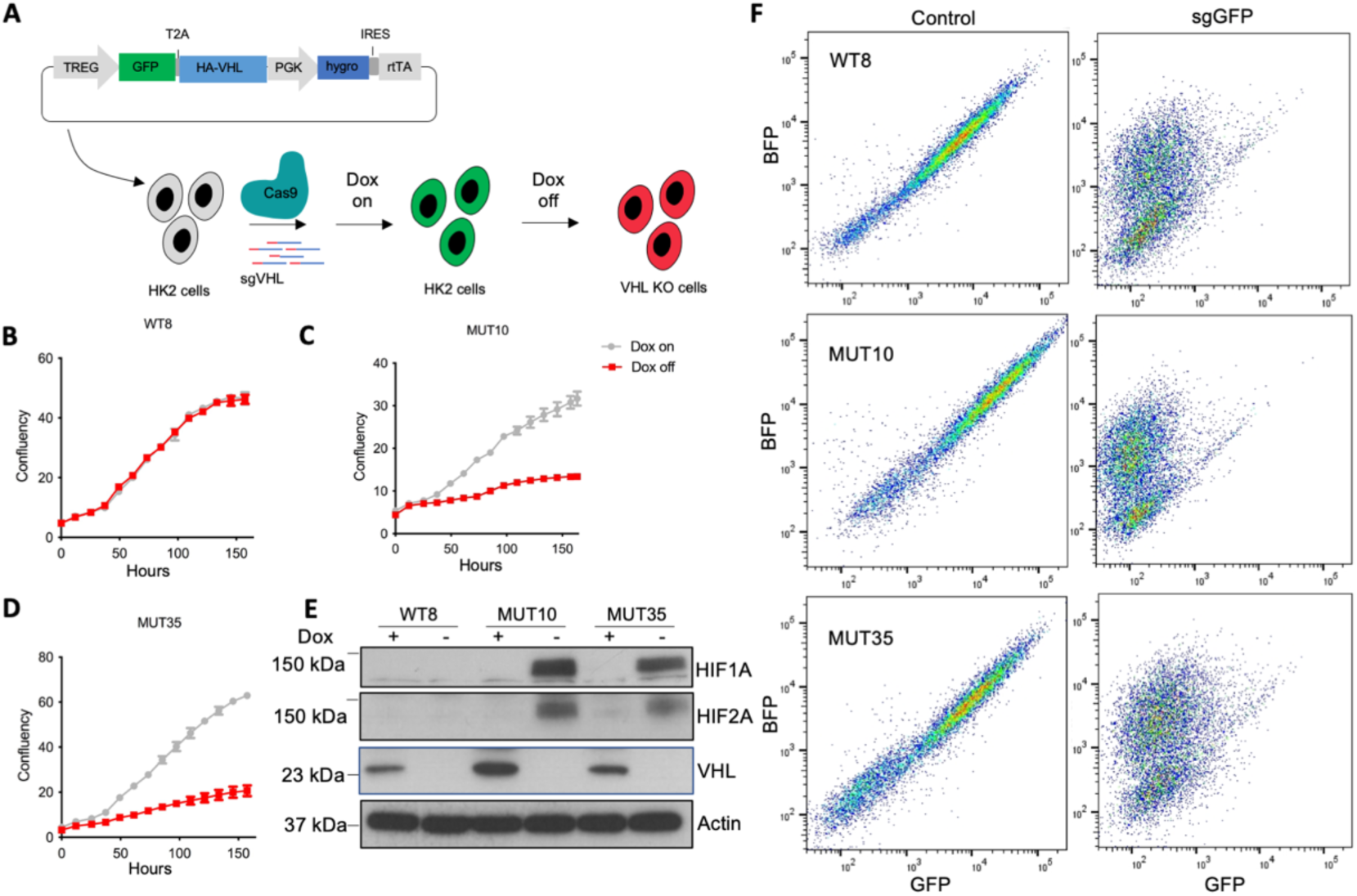
Establishment of models with doxycycline-controllable VHL expression. **A.** Schematic of doxycycline (dox) inducible VHL re-introduction into VHL mutant (MUT10 and MUT35) and wildtype control (WT8) clones. **B-D.** Proliferation of WT8 (B), MUT10 (C) and MUT35 (D) cells with and without dox. N=2 replicates per condition. (Mean and S.E.M). **E.** Western blot of HIF1A, HIF2A, VHL and Actin on WT8, MUT10 and MUT35 cells with and without dox. **F.** Cas9 editing efficiency tested on WT8, MUT10 and MUT35 cells by a reporter plasmid using fluorescence-activated cell sorting.

**Fig. S3.**
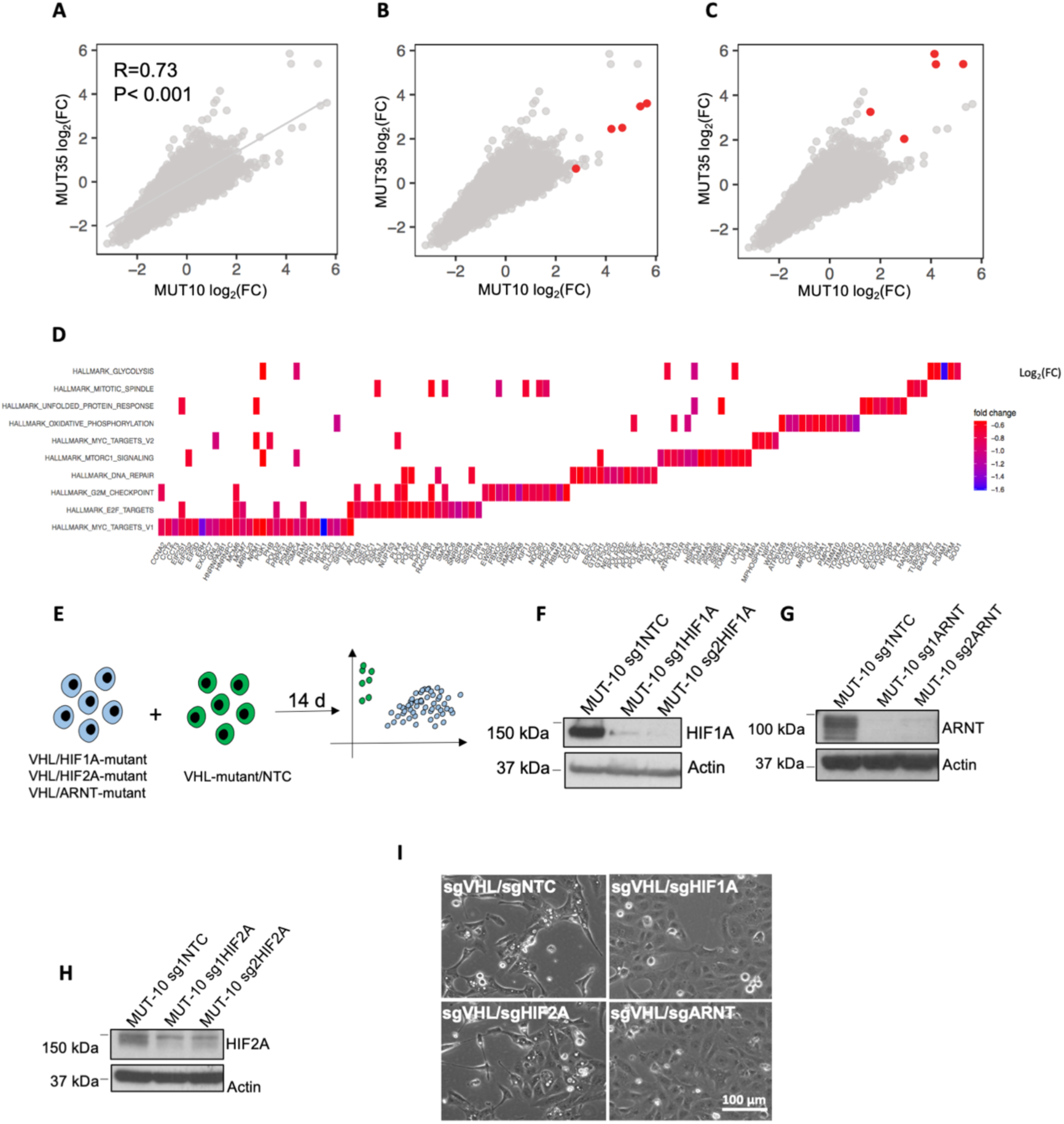
HIF1A and ARNT inhibit proliferation upon VHL inactivation. **A.** CRISPR/Cas9-based genome wide screen data. sgRNA abundance on day 28 relative to start of the assay in VHL mutant clones MUT10 and MUT35. R, Pearson’s correlation coefficient. **B-C.** As in A with sgRNAs targeting genes of interest highlighted in red: HIF1A in B and ARNT in C. **D**. Pathway enrichment analysis on the top 500 genes the sgRNAs of which are depleted over time in MUT10 and MUT35 cells using the Cancer Hallmarks gene sets. **E.** A schematic of the competitive proliferation assay. VHL-HIF1A, VHL-HIF2A and VHL-ARNT double mutant cells (BFP labelled) competed against VHL-NTC single mutant cells (GFP labelled). **F-H.** Western blot of HIF1A, ARNT and HIF2A on MUT10 cells with and without HIF1A, ARNT or HIF2A inactivation, respectively. **I**. Morphology of VHL-NTC, VHL-HIF1A, VHL-HIF2A and VHL-ARNT cells.

**Fig. S4.**
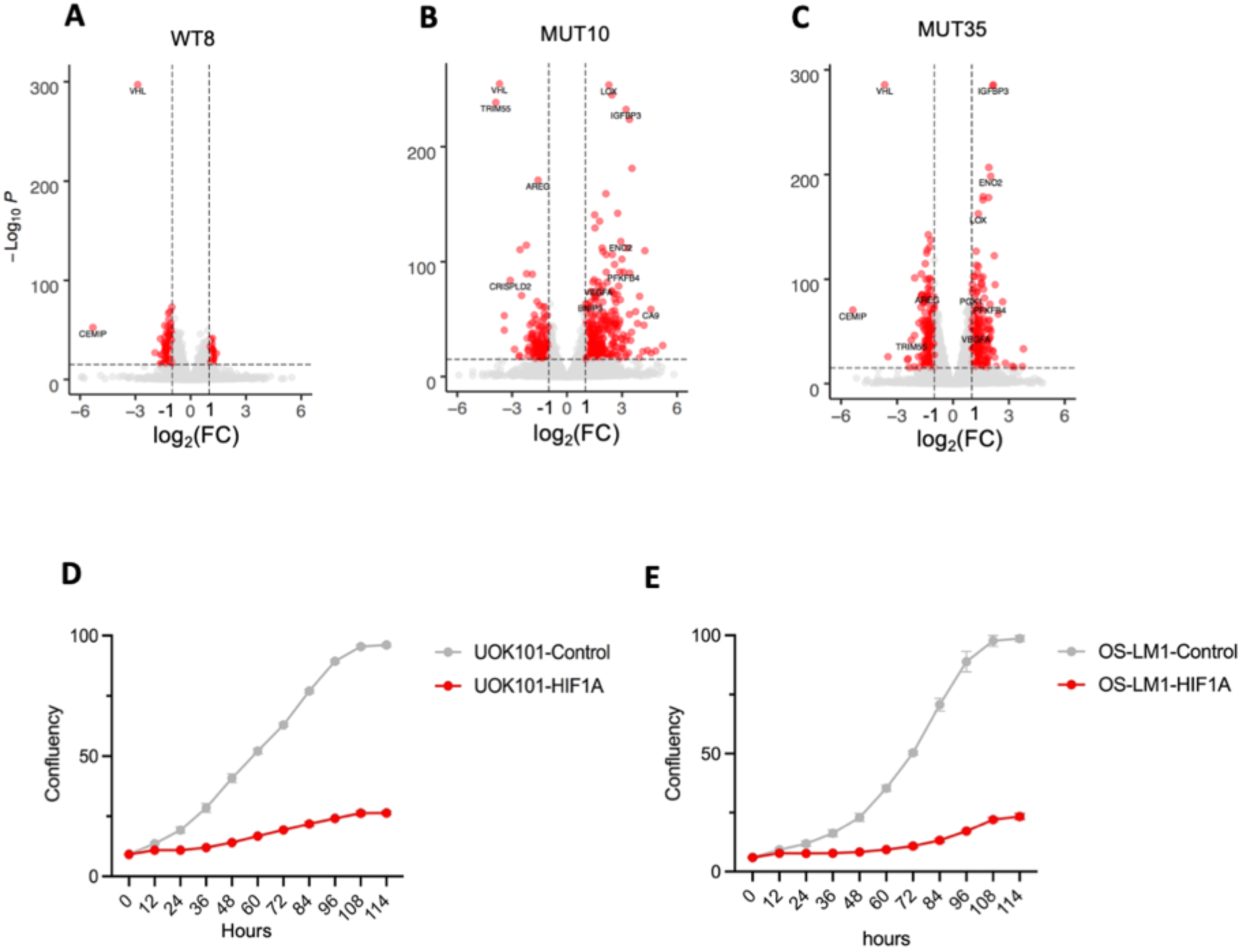
HIF1A inhibits mitochondrial function. **A-C.** Differential gene expression analysis by RNA-seq. WT8, MUT10 and MUT35 cells, dox off compared to dox on. Adjusted p-value and fold change determined by DESeq2. **D.** Proliferation of UOK101 ccRCC cells with and without HIF1A cDNA expression. N=2 replicates per condition (Mean and S.E.M). **E.** Proliferation of OS-LM1 ccRCC cells with and without HIF1A cDNA expression. N=2 replicates per condition (Mean and S.E.M).

**Fig. S5.**
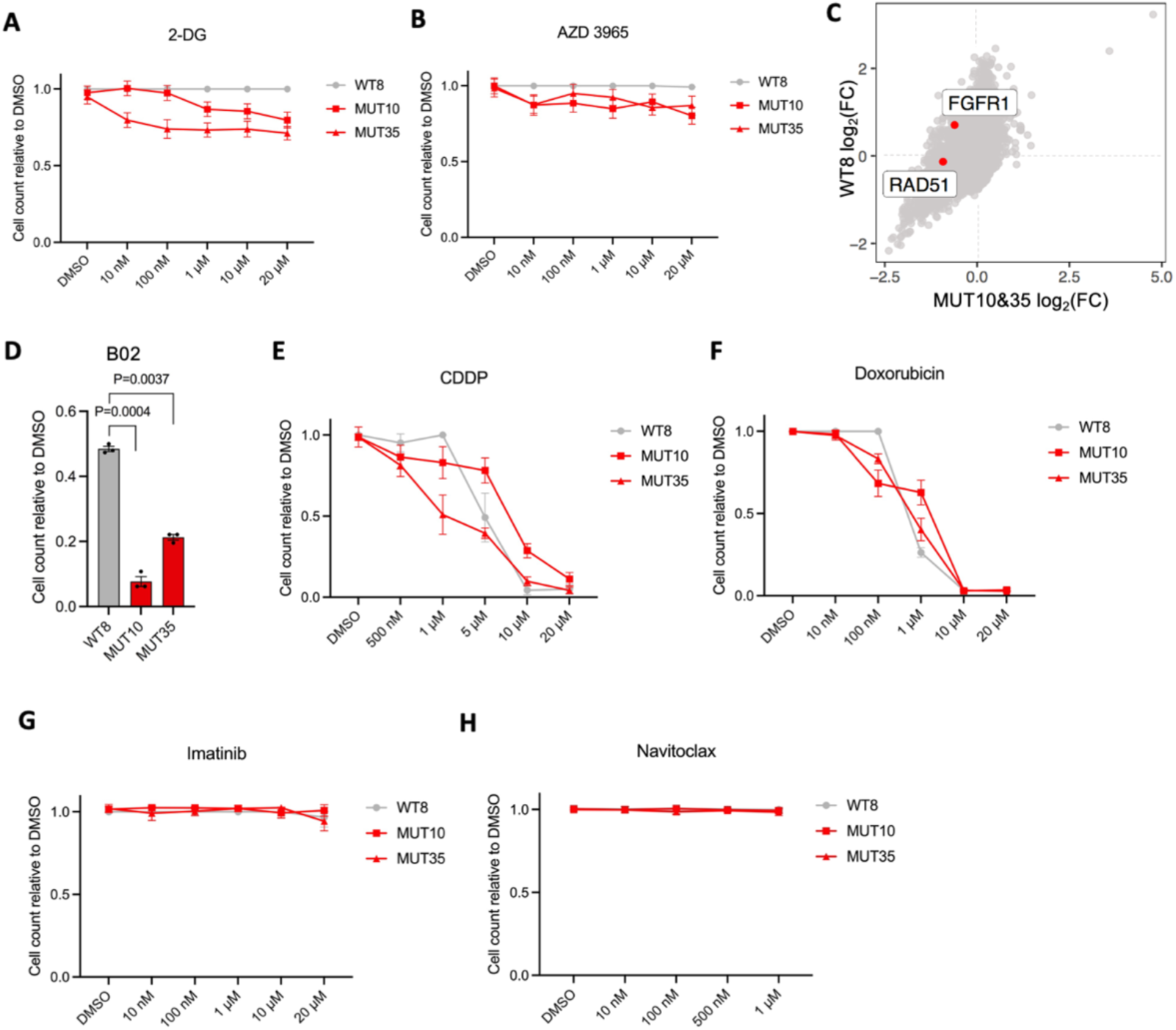
*VHL* mutant cells show druggable genetic vulnerabilities. **A-B.** Cell counts on day 5 relative to DMSO control group under 2-DG (10 nM, 100 nM, 1 μM, 10 μM, 20 μM) and AZD3965 (10 nM, 100 nM, 1 μM, 10 μM, 20 μM) treatments. N=4 per condition (Mean and SD). **C.** CRISPR-Cas9 screen data. FGFR1and RAD51 gene beta score distribution in in VHL WT and mutant cells. **D.** Cell counts on day 5 relative to DMSO control group under B02 (10 μM). N=3 replicates per condition (Mean and SD). Paired t test. **E-H**. Cell counts on day 5 relative to DMSO control group under CDDP (10 nM, 100 nM, 1 μM, 10 μM, 20 μM), Doxorubicin (10 nM, 100 nM, 1 μM, 10 μM, 20 μM), Imatinib (10 nM, 100 nM, 1 μM, 10 μM, 20 μM) and Navitoclax (10 nM, 100 nM, 1 μM, 10 μM, 20 μM).

**Fig. S6.**
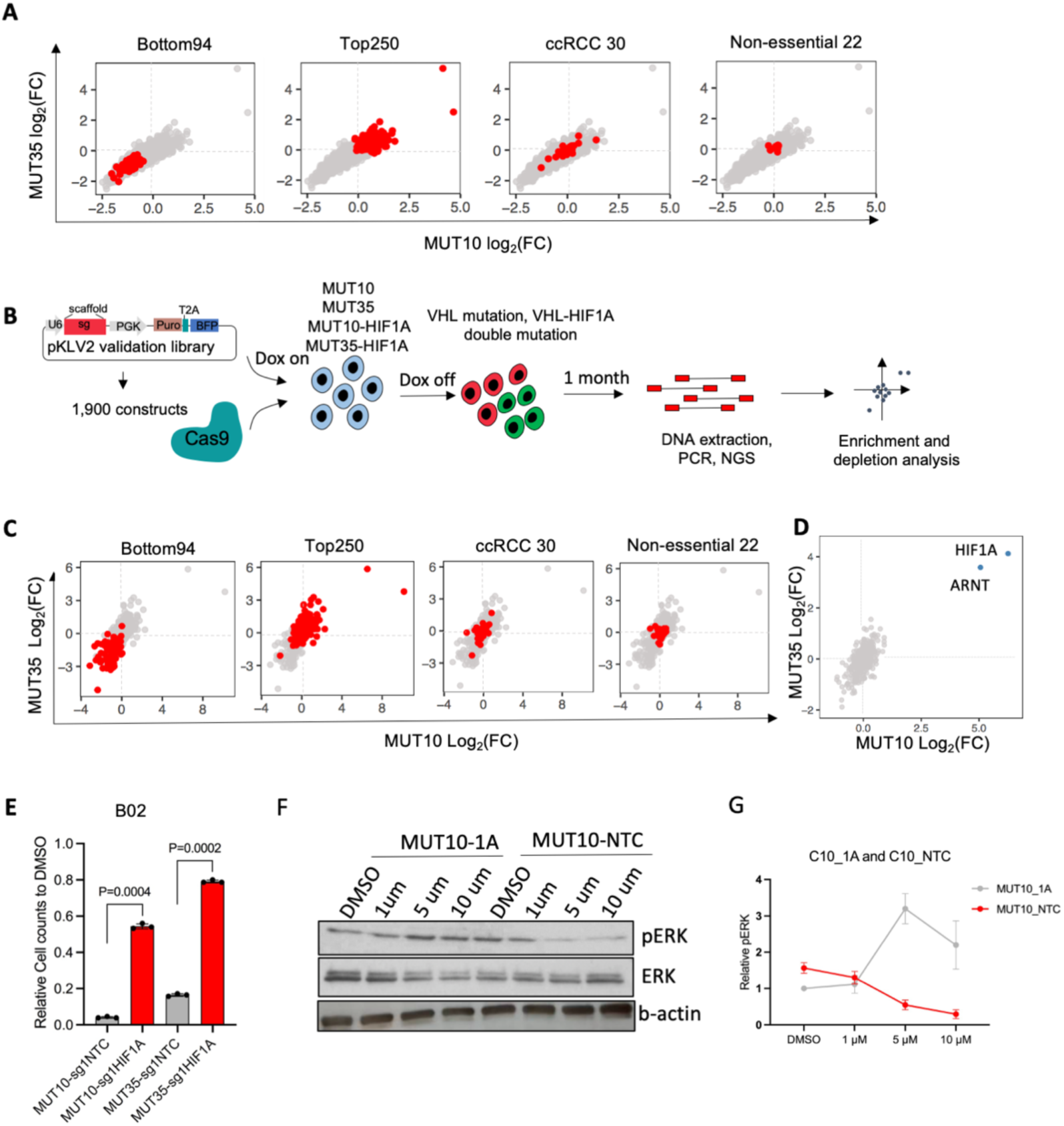
CRISPR-Cas9 validation screen. **A.** Genes selected for the validation screen. Distribution of beta scores in the genome-wide CRISPR-Cas9 screen data set. 94 genes the constructs of which were specifically depleted in *VHL* mutant cells, 250 the constructs of which were specifically enriched in *VHL* mutant cells, 30 frequently mutated ccRCC genes and 22 non-essential control genes. **B.** Schematic of the validation screen on MUT10, MUT35, MUT10-sgHIF1A and MUT35-sgHIF1A cells. **C.** CRISPR-Cas9-based validation screen data. Gene level construct abundance relative to start of the assay in MUT10 and MUT35 cells. Gene sets of interest highlighted in red. **D.** CRISPR-Cas9-based validation screen data, doxycycline withdrawal before sgRNA transduction. Gene level construct abundance relative to start of the assay in MUT10 and MUT35 cells. **E.** Cell counts of B02 (10 μM) treated cells on day 5 relative to DMSO control group. N=3 replicates per condition (Mean and SD). Student’s t-test. **F**. WB of MUT1-1A and MUT10-NTC treated with DMSO, 1μM, 5μM,10 μM Pazopanib for pERK, total ERK and beta-actin. **G**. Quantification of relative pERK at different treatment concentration at indicated cell population.

**Fig. S7.**
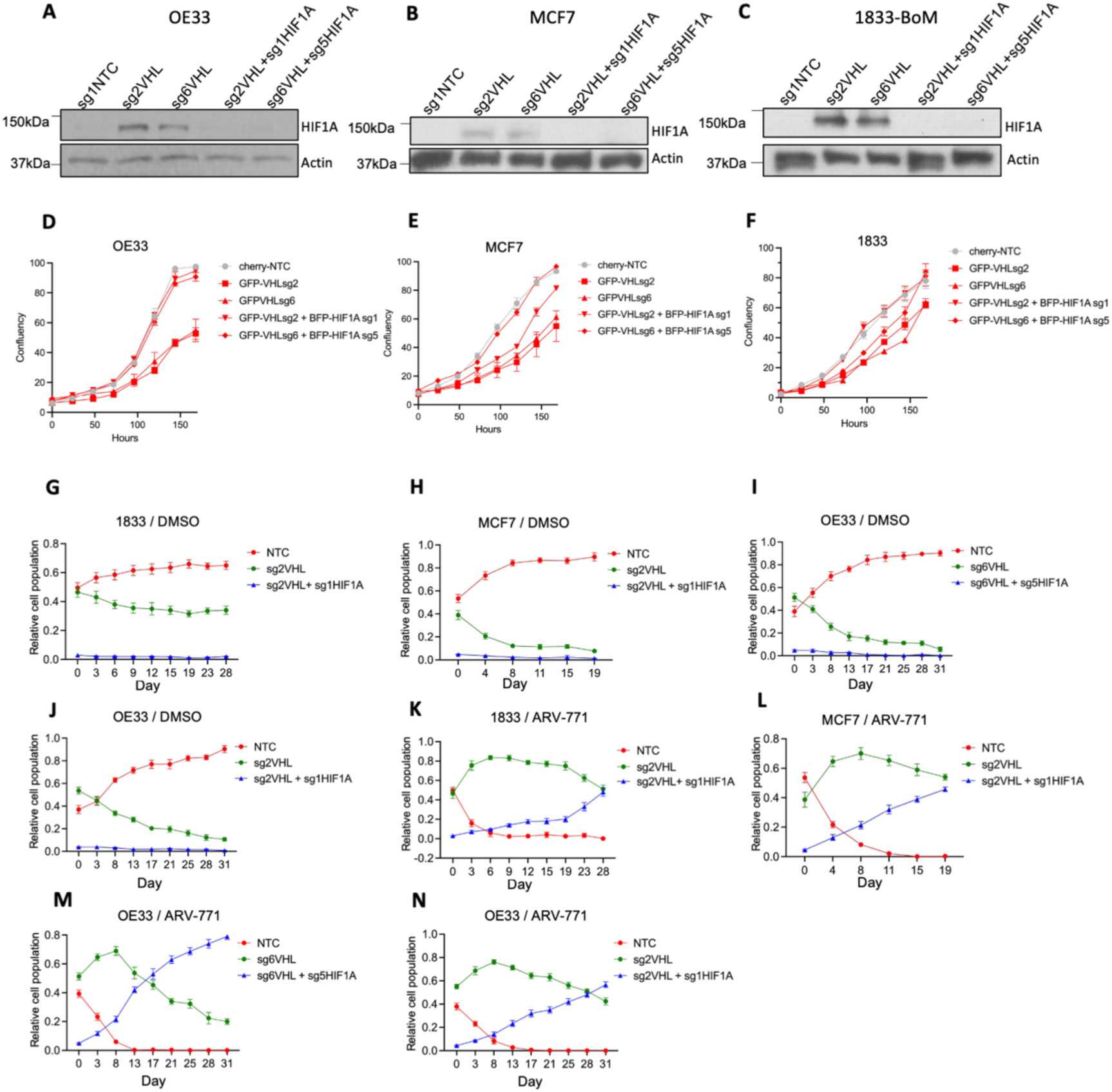
HIF1A loss-mediated escape from PROTAC-induced growth inhibition in cancer. **A-C.** Western blot of HIF1A and Actin on WT (NTC), VHL mutant (VHLsg2, VHLsg6) and VHL-HIF1A double mutant (VHLsg2/HIF1Asg1, VHLsg6/HIF1Asg5) OE33 (A), MCF7 (B) and 1833-BoM (C) cells. **D-F**. Incucytes proliferation of OE33, MCF7 and 1833 under different genotype. N=3. **G-N.** FACS-based quantification of the relative abundances of different cell populations in competition assays. 1833-BoM cells: 500 nM ARV-771; MCF7 cells: 200 nM ARV-771; OE33 cells: 400 nM ARV-771. N=3 for each condition and timepoint (Mean and SD).

